# A single TRPV1 amino acid controls species sensitivity to capsaicin

**DOI:** 10.1101/732701

**Authors:** Ying Chu, Bruce E. Cohen, Huai-hu Chuang

## Abstract

Chili peppers produce capsaicin, the principle chemical that accounts for the culinary sensations of heat. Capsaicin activates the transient receptor potential cation channel subfamily V member 1 (TRPV1) on sensory neurons to alter the membrane potential to induce pain. While structural studies have identified residues important for the binding of capsaicin to rat TRPV1, there is still no clear understanding of differential capsaicin sensitivity of TRPV1 between birds and mammals. To determine the residues dictating relative capsaicin sensitivity among species, we have used intracellular Ca^2+^ imaging to characterize chimeras composed of capsaicin-sensitive rat TRPV1 (rTRPV1) and capsaicin-insensitive chicken TRPV1 (cTRPV1) with a series of capsaicinoids. We find that chimeras containing rat E570-V686 swapped into chicken receptors displays capsaicin sensitivity, and that a single amino acid substitution in the S4-S5 helix, changing the alanine at position 578 in the chick receptor to a glutamate, is sufficient to endow micromolar capsaicin sensitivity. Moreover, introduction of lysine, glutamine or proline at A578 also install capsaicin sensitivity in cTRPV1. Comparing the derivatives Cap-EA and Cap-EMA with capsaicin, these two compounds with the hydrophilic vanilloid-like moiety affect the protein-ligand interaction. The ability of 10 μM Cap-EA to activate cTRPV1-A578E and the differential response of mutants to the analogs suggests that chick A578 may participate in vanilloid binding, as does the corresponding rTRPV1 site. The hydrophilic vanilloid agonist zingerone 500 μM failed to activate any A578 mutants that retained capsaicin sensitivity, suggesting that the vanilloid group alone is not sufficient for receptor activation. Replacing the rTRPV1-E570 residue with K, Q shows a similar tendency to maintain the receptor capsaicin sensitivity. Our study demonstrates a subtle modification on different species TRPV1 globally alter their capsaicin response.

## Introduction

Pain occurs following intense or damaging stimulation of nociceptors, peripheral sensory neurons that express the transient receptor potential cation channel subfamily V member 1 (TRPV1). TRPV1 functions as integrator of physical and chemical noxious stimuli. When activated by noxious chemicals, heat (>43°C), or acid (pH≦ 5.9), TRPV1 permits cation influx to elicit membrane currents (Caterina et al., 1997; Tominaga et al., 1998; Wood et al., 1988). TRPV1 allows these primary afferent neurons to distinguish noxious environmental signals from innocuous events, protecting organisms against further injury. A key TRPV1 ligand is capsaicin, a small lipophilic molecule from chili peppers that induces sensations of heat. Capsaicin and protons each lower the temperature threshold of TRPV1 activation, so that it is activated at room temperature (Jordt et al., 2000).

Besides capsaicin, TRPV1 may also be activated by plant-derived pungent agents, particuarly vanilloid-containing compounds, such as resinfetrontoxin (Raisinghani et al., 2005), piperine (McNamara et al., 2005), and zingerone (Liu et al., 2000; Szallasi and Blumberg, 1999). TRPV1 channels cloned from different species share conserved function as noxious stimuli integrators, but their sensitivity to pungent chemicals varies among the orthologues. Rodent and human TRPV1 channels have high vanilloid sensitivities, with nanomolar EC_50_ value (Correll et al., 2004; Hayes et al., 2000; Savidge et al., 2002). Frog and rabbit TRPV1 channels are less sensitivite to capsaicin, (Gavva et al., 2004; Ohkita et al., 2012) and the chicken TRPV1 shows the least sensitivity (Jordt and Julius, 2002). The pharmacological characteristics of TRPV1 provides the understanding to efficiently target, activate and inhibit the channel, which may be one of the strategies to design analgesic drugs (Brito et al., 2014; Nersesyan et al., 2017; Szallasi et al., 2007).

Studies with synthetic water-soluble capsaicin analogs (Jung et al., 1999) and mutational analysis highlights critical residues in intracellular domain of S1-S4 segments in charge of capsaicin sensitivity. By constructing chimeric receptors between capsaicin-sensitive rat TRPV1 (rTRPV1) and capsaicin resistant chicken TRPV1 (cTRPV1), Jordt discovered specific mutations of the Y511 or S512 residue specifically suppress the capsaicin responsiveness without affecting proton-induced current of the rat receptors (Jordt and Julius, 2002). For rabbit TRPV1 (oTRPV1), mutating I550 and L547 (the equivalent residue for T550 and M547 in the rat receptor) enhances capsaicin sensitivity and resinferotoxin (RTX) binding respectively (Chou et al., 2004; Gavva et al., 2004). The A561 and Y523 (the T550 and Y512 in the rat receptor) were reported as the reason for the limited capsaicin sensitivity of frog TRPV1 (xTRPV1) (Ohkita et al., 2012).

Being permeable to the plasma membrane, capsaicin binds to the intracellular face of TRPV1 (Jung et al., 1999) and was reported to pass into the cytosol to activate TRPV1 channels on the ER (Gallego-Sandin et al., 2009). Cryo-EM structural studies of TRPV1 (Cao et al., 2013; Gao et al., 2016) show that E570 of rTRPV1 is close to the interface of cytosol and protein. E570 and Y511 constitute the bottom part of the ligand binding pocket which contains RTX ligand density, including its vanilloid headgroup (Cao et al., 2013; Gao et al., 2016). Capsaicin has been described to bind in an “tail-up, head-down” orientation in a vanilloid-binding pocket, with the vanillyl head group facing the residues closest to the cytosol (Yang et al., 2015).

Despite this structural evidence for ligand-TRPV1 interactions, questions remain about how capsaicin activates the channel and why TRPV1 channels from different species show radically different capsaicin response. In this study, we reveal the molecular mechanisms of capsaicin sensitivity differences between chicken and rat TRPV1. We express TRPV1 channels in HEK293T cells and quantify its activation by ratiometric Ca^2+^ imaging to identify specific amino acids responsible for ligand activation. These mutagenesis studies show that changing a single residue, A578 in the chick receptor, is sufficient to confer micromolar capsaicin sensitivity on cTRPV1.

## Methods and materials

### Molecular cloning

Wild-type rat (*Rattus norvegicus*) and chicken (*Gallus gallus*) TRPV1 genes on pcDNA3 plasmid were used to construct chimeras by overlap extension PCR, to swap in the rTRPV1 sequence to cTRPV1 corresponding sites, including rTRPV1 N-terminus (M1-R428; named as Ch6), S1-S4 (F429-I569; named as Ch3/12), S5-S6 (E570-V686; named as Ch9-18), S1-S6 (F429-V686; named as Ch3/18), and C-terminus (N687-K838; named as Ch15). For single point mutated cTRPV1 and rTRPV1, the genes were cloned into pxpIV plasmids and linked with three repeats of HA tag (3XHA) at the N-terminus for Western blotting and immunostaining. Point mutations were introduced by QuikChange with PfuUltra II Fusion HS DNA Polymerase (Aligent). To remove rTRPV1-G602-N625 (GKNNSLPMESTPHKCRGSACKPGN) sequences from Ch9/18 and rTRPV1 genes, the sequence was deleted by back-to-back PCR (Phusion Hot Start Flex DNA Polymerase, New England Biolab) and ligated to blunt ends (T4 DNA ligase, Thermo Scientific). Plasmids were sequenced by Genomics BioSci & Tech, and then transformed and amplified in DH5α competent cells (Yeastern Biotech).

### Mammalian cell culture

HEK293T cells were grown in MEM/EBSS (HyClone) medium with 9% fetal bovine serum (FBS, Gibco), and 100 U/ml penicillin and 100 μg/ml streptomycin (Lonza). The incubator was kept at 37℃ with 5% CO_2_. The cells were seeded onto plates one day before transfection, and reached 60-90% confluency on the time for transfection. OptiMEM (Life Technology) and Avalanche®-Omni Transfection Reagent (EZ Biosystems) were mixed with plasmids and added into wells with HEK293T cells. After two days, the transfected cells were prepared for Ca^2+^ imaging, immunostaining or Western blotting.

### Ratiometric Ca^2+^ imaging

The 96-well plates were coated with poly-D-lysin (0.1 mg/ml) and collagen (55 μg/ml). The transfected cells were added to 96-well plates with MEM + 5.4% FBS + penicillin/streptomycin and grow overnight. Cells were loaded with 0.02% pluronic F-127 (Life Technology), and 2 μM Fura-2 AM (Life Technology) (Grynkiewicz et al., 1985; Yates et al., 1992) for 3-5 hours in imaging solution [8.5 mM HEPES, 140 mM NaCl, 3.4 mM KCl, 1.7 mM MgCl_2_, and 1 mM CaCl_2_, pH 7.4] at 30℃ with 5% CO_2_. Solutions were replaced with the same imaging solution without Fura-2 AM before imaging. Background-subtracted, emitted fluorescence following excitation at 340 nm and 380 nm were detected by an EMCCD camera (Photometrics, Evolve) driven by Slidebook 6 digital microscopy software (Intelligent Imaging Innovations). Fluorescence data were acquired by capturing the frame rate at one frame every 5 sec with 20-50 ms exposure time to either wavelength. All Ca^2+^ imaging experiments were conducted at 22° C.

Capsaicin (Pfaltz & Bauer), zingerone (Pfaltz & Bauer), and cocktail were prepared as stock solutions in DMSO (Calbiochem). The cocktail solution used to induce maximum TRPV1 activation contained capsaicin 100 μM, resinferatoxin (Ascent) 5 μM, phenylarsine oxide (Alfa Aesar) 100 μM. The cocktail ligands were dissolved in solution that replace 140 mM NaCl with 140 mM CsCl. Reactive chemical or ligands were prepared as 2X concentration stocks (2-fold).

Concentration of ligands reached 1X after applying to wells containing equivalent amount of imaging solution (75 μl). The CsCl concentration in cocktail solution was 70 mM after addition to imaging solutions. For pre-treatment experiments, dye-loaded cells were incubated with 1X concentration chemicals in imaging solution without Ca^2+^. 1X concentration ligand with 2 mM CaCl_2_ was added during recording.

The agonists were added at the 5^th^ frame after the start of recording. The background 340/380 ratio value was calculated as the average ratio of 0^th^-4^th^ frames of all cells. The agonist-induced activation was calculated as average maximum 340/380 ratios of each cell between 5^th^-34^th^ frames after substracting the background value.

### Western blotting

Transient transfected HEK293T cells were homogenized in lysis buffer [150 mM NaCl, 50 mM Tris pH 7.5, 2 mM EDTA, 0.5% Triton X-100, 0.5% NP-40] mixed with protease inhibitor cocktail set III (EDTA-free, Calbiochem). Concentrations of protein samples were measured by BCA assay (Thermo Scientific, Pierce). Protein lysates 25 μg mixed with 6Xsample buffer with 1 mM DTT was resolved in SDS-PAGE and transferred onto PVDF. HA-tagged TRPV1 protein was identified by rabbit polyclonal primary antibody, HA.11 (Covance PRB-101P, 1:2000 dilution). GAPDH was detected by rabbit polyclonal GAPDH antibody (Santa Cruz Biotechnology FL-335, 1:5000 dilution). Both antibodies were visualized by anti-rabbit IgG HRP conjugated secondary antibody (Pierce prod#31430, 1:5000 dilution) and the blot images were acquired using BioSpectrum 810 (UVP). The images were analyzed with ImageJ. The HRP signal of each band were normalized to the rTRPV1 band on the same membrane in order to compare the results on different day and different membrane.

### Immunostaining and microscopy imaging

Chamber slides were coated with PDL and collagen for overnight separately. The transient transfected HEK293T cells were seeded onto PDL and collagen-coated chamber slides and incubated at 37 ℃, 5% CO_2_ overnight. The cells were fixed with 4% paraformaldehyde in 1X DPBS, and permeabilized with 1X DPBS + 0.5% Triton X-100. After treatment with blocking buffer [1X DPBS + 1% bovine serum albumin (BSA, *w*/*w*) + 0.5%Triton X-100] for 1 hour, the sample was incubated with primary and secondary antibody, HA.11 Clone 16B12 monoclonal antibody (Covance MMS-101P, 1:1000) and Alexa Fluor 488 goat anti-mouse Ab (Life Technology A11001, 1:5000) respectively. The slides were stained by DAPI, mounted in SlowFade Diamond antifade (Life Technology), and sealed with coverslip. Images were acquired with LSM780 laser scanning confocal microscope (Carl Zeiss).

### Capsaicinoid synthesis

*O*-aminoethyl capsaicin (**cap-EA**) was synthesized in a 2-step procedure from capsaicin. (ref: Li, PNAS 108, 8497). Boc-aminoethyl bromide was dissolved in 3 mL of dry THF and added to capsaicin (305 mg, 1.0 mmol), potassium *t*-butoxide (112 mg, 1.0 mmol), and 18-crown-6 (264 mg, 1.0 mmol) dissolved in 7 mL of dry THF. The reaction was stirred overnight, concentrated, and partitioned between EtOAc and saturated NH_4_Cl solution. The organics were washed well with water, then dried and concentrated to 439 mg of crude white solid. Of this, 150 mg (0.33 mmol) was dissolved in 1 mL of CH_2_Cl_2_, and 35 μL (0.35 mmol) of thiophenol and 0.5 mL of TFA were added. After 30 min, the solvent was evaporated and the residue purified by HPLC on a semi-preparative 18 column, eluting with a linear gradient of 20 to 70% CH_3_CN in 50 mM NH_4_OAc, pH 4.5, over 50 min. The cap-EA peak eluted at 40 min (52% CH_3_CN) and free capsaicin at 55 min (65% CH_3_CN). Fractions were lyophilized to a white powder (81 mg, 54%). ^1^H NMR (DMSO-*d*_6_, 400 MHz) showed additional peaks relative to capsaicin at: 4.66 (t, 2H) and 3.39 (t, 2H). MS calcd for C_20_H_33_N_2_O_3_+ (MH^+^): 349.2. Found: 349.2.

*O*-(*N*-methylaminoethyl) capsaicin (**cap-EMA**): the acetate salt of cap-EA (20 mg, 49 *μ*mol) was dissolved in 1 mL of dry DMF with triethylamine (7 *μ*L, 50 *μ*mol) and cooled to 4° C. Iodomethane (2 *μ*L, 33 *μ*mol) was dissolved separately in 1 mL of dry DMF and added dropwise to the reaction over 10 min. The reaction was stirred for 2 h at 4° C, and then purified by reverse phase HPLC as above, with product eluting at 43 min (55% CH_3_CN), The fractions were lyophilized to white powders. By ^1^H NMR (DMSO-*d*_6_, 400 MHz), cap-EMA showed additional peaks relative to capsaicin at: 4.62 (t, 2H), 3.38 (t, 2H), and 2.90 (s, 3H). MS calcd for C_21_H_35_N_2_O_3_^+^ (MH^+^): 363.5. Found: 363.4.

## Results

### Chimeras with rTRVP1 S5-S6 have similar capsaicin sensitivity as those with rTRPV1 S1-S4

TRPV1 forms homotetrameric channels, with each subunit consisting of six α-helical transmembrane domains connected by two intracellular N- and C-terminal segments. Transmembrane domain 5 and 6 (S5-S6) constitutes the central pore separated by an intervening pore loop that presumes the ion selective filter. Transmembrane segments S1-S4 surrounding the central pore remain static, while the S4-S5 linker moves to prompt TRPV1 opening (Cao et al., 2013; Liao et al., 2013). To identify residue(s) critical for capsaicin sensitivity in TRPV1, we firstly analyzed rat-chicken TRPV1 chimeras (Fig. 1A. and Materials and Methods) to narrow down regions important for capsaicin sensitivity of rat. We used ratiometric Ca^2+^ imaging (O’Connor and Silver, 2013) with Fura-2 to quantify the intracellular Ca^2+^ raise due to TRPV1 activation. Capsaicin elicits Ca^2+^ signals in rTPPV1 transfected HEK293T cells. To verify the expression and function of mutated TRPV1 responding weakly to capsaicin, a powerful agonist cocktail solution was applied to stimulate maximum channel activation. The cocktail solution contains 100 μM capsaicin, 5 μM resinferatoxin (RTX), 100 μM phenylarsine oxide (PAO) and 70 mM cesium ion. Besides two strong vanilloids, the capsaicin and RTX (Raisinghani et al., 2005; Szallasi et al., 1999), PAO functions as a surrogate for oxidative stress that potentiate activation of chicken and rat TRPV1 (Chuang and Lin, 2009), while cesium ion maximizes the responses. Cocktail solution efficiently activates both rTRPV1 and cTRPV1, eliciting extracellular Ca^2+^ entry that can be blocked by 1 μM ruthenium red (Dray et al., 1990) (Fig. 1B). Ch3/18 with six transmembrane domains acquires highest capsaicin sensitivity among chimeras. The EC_50_ representing by 340/380 ratio was slightly higher than wildtype rTRPV1 but still within several hundred nanomolar. Dividing rTRPV1 transmembrane domains into two regions generates two capsaicin-sensitive chimeras, Ch3/12 and Ch9/18 with EC_50_ in several micromolar (Fig. 1C). N-terminus swaps provide limited improvement, while rTRPV1 C-terminus (Cap 340/380 ratio=0.02±0.003, N=3) inhibits the chimeras with cTRPV1 backbone (Fig. 1D). The residues Y511, S512 (Jordt and Julius, 2002), M547, and T550 (Gavva et al., 2004; Ohkita et al., 2012) reported to contribute capsaicin sensitivity are in the S1-S4 that possessed by Ch3/12. We further confirm the residue dictating capsaicin sensitivity in Ch9/18.

**Fig 1.**
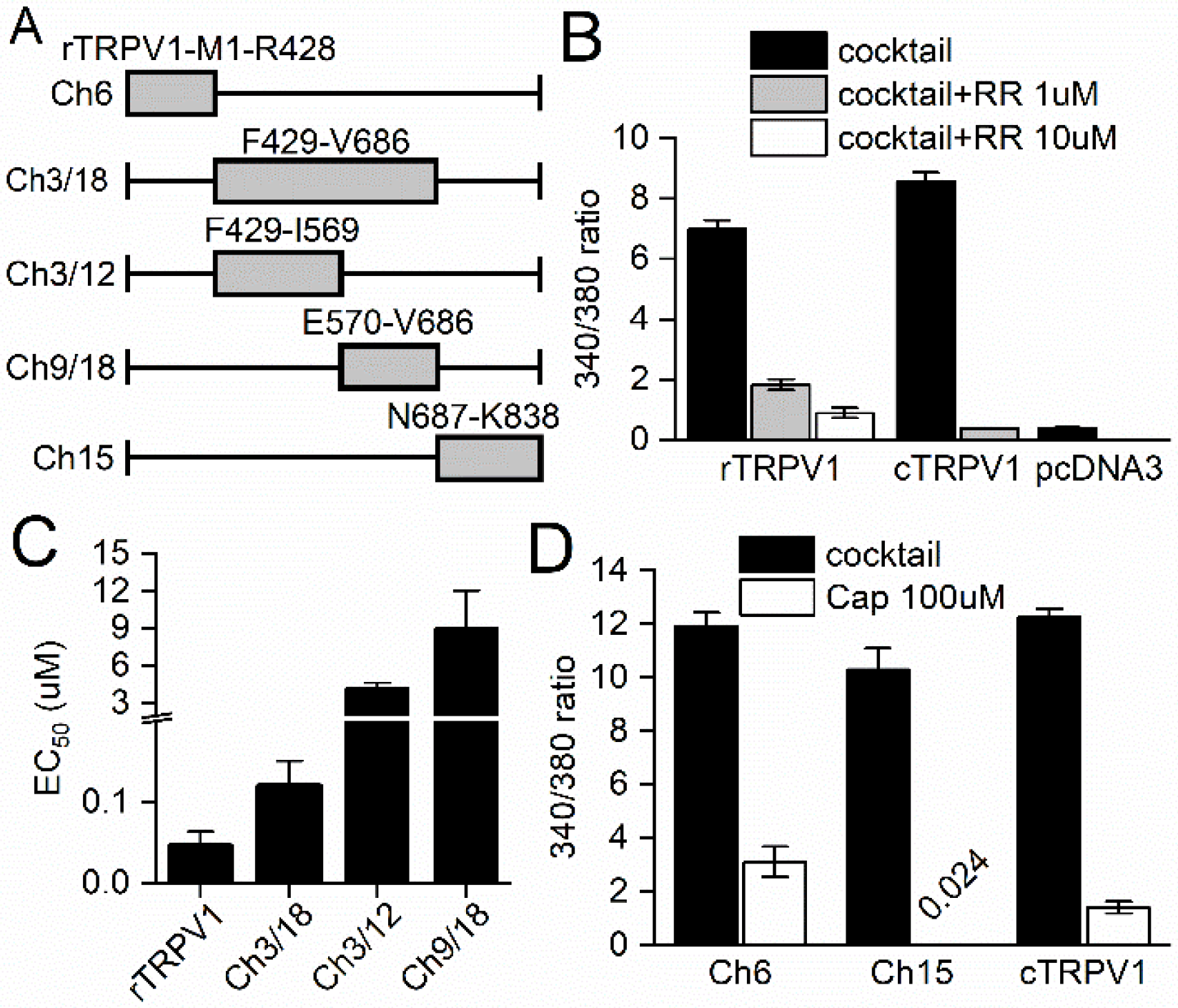
Comparing rat/chicken TRPV1 chimeric protein capsaicin sensitivity. (A) The boundaries between chicken and rat TRPV1 of chimeric proteins. (B) Quantitative data for the activation and inhibition of rat and chicken TRPV1 with cocktail and ruthenium red (N=3∼4). (C) Different concentration capsaicin induced activation on capsaicin sensitive chimeras (N=3). (D) Response of chimeras with rat N or C-terminus to capsaicin and cocktail solution (N=3).

### A single mutation is sufficient to transfer capsaicin sensitivity to cTRPV1

The residues that make Ch9/18 different from cTRPV1 can be classified into nine parts: E570A, F589L, S632Y, D654R, A657S, I660V, L664V, A665L (rat/chicken equivalent residue), and rTRPV1-G602-N625, a region with high sequence divergence. Point mutagenesis or deletion of rTRPV1-G602-N625 segment was performed on Ch9/18 to identify the residue(s) that reduce capsaicin sensitivity. The equivalent sites on rTRPV1 are shown in Fig. 2A. 100 μM capsaicin-induced activation 340/380 ratio values were adjusted by dividing to cocktail induced ratio increase to eliminate effect of non-specific loss of channel function. Ch9/18-E570A loses most of the capsaicin sensitivity. Deleting G602-N625 decreases capsaicin sensitivity of Ch9/18 and the cocktail induced maximum channel opening. However, deleting the region on wildtype rTRPV1 decrease both the capsaicin and cocktail induced activation. G602-N625 region is non-essential to specifically affect the capsaicin sensitivity (Fig. 2B).

**Fig 2.**
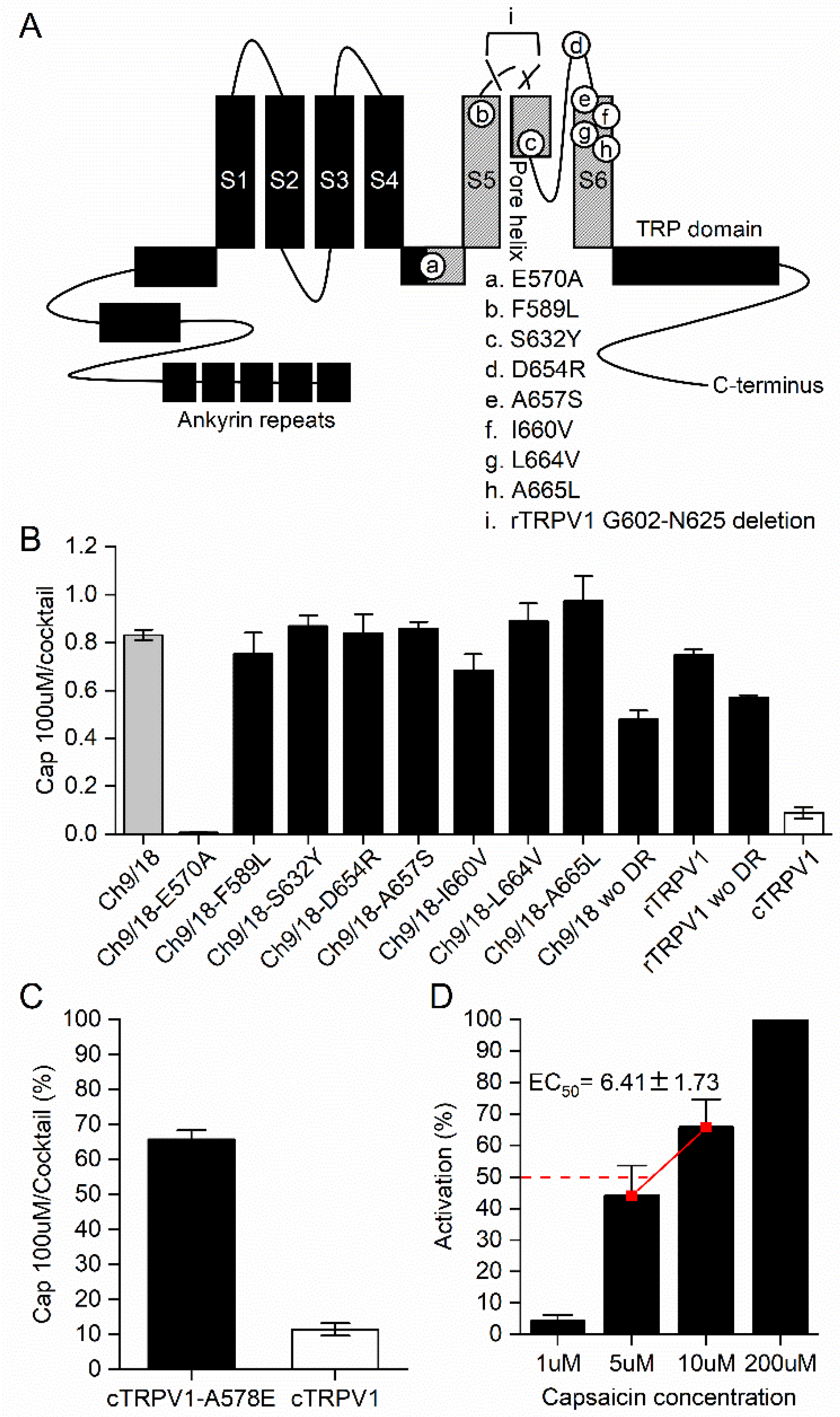
Identify the residue endow capsaicin sensitivity on cTRPV1. (A) Mutagenesis sites on Ch9/18. (B) The ratio of capsaicin 100 μM (N=3) and cocktail (N=2∼3) induced response represent the capsaicin specific activation of mutant. (C) Comparing the cTRPV1-A578E to cTRPV1 100 μM capsaicin-cocktail responsiveness (N=3). (D) Determine the EC_50_ of cTRPV1-A578E (N=3).

A578 on cTRPV1 is mutated to glutamic acid to verify the importance of E570, the equivalent residue in rTRPV1, for capsaicin sensitivity. A578E mutation successfully restores capsaicin sensitivity onto cTRPV1 (Fig. 2C) with EC_50_ at micromolar (Fig. 2D). The E570 had been predicted to participate in human and rat TRPV1 vanilloid pocket formation through computational modeling. Mutational experiment on E570 (E571 in the paper) present an impaired function of TRPV1 to open (Lee et al., 2011; Yang et al., 2015).

### Multiple amino acid replacements on cTRPV1-A578 possess capsaicin sensitivity

To further elucidate the mechanism by which the glutamic acid introduces capsaicin sensitivity to cTRPV1, we systematically replaced the A578 on cTRPV1 and E570 on rTRPV1 to the other 19 amino acids. The A578E mutant doesn’t show higher expression level or membrane distribution than the non-sensitive mutants in results of Western blotting (Fig. 3A) and imaging of protein distribution (Fig. 3B) determined with HA-tag antibodies. The results suggest that the residue specifically increase capsaicin sensitivity. Capsaicin 30 μM were used as standard to determine capsaicin-sensitive mutants in cTRPV1 mutants. The cocktail efficiently activates most cTRPV1-A578 mutants (Table S1) except A578D, F, G, and W which intracellular 340/380 ratio increment are lower than six. The response of cTRPV1-A578 mutant to capsaicin was adjusted to cocktail induced maximum channel activation. The A578K, Q, and P are the most efficient choices to endow capsaicin-sensitivity to cTRPV1 other than glutamic acid (Fig. 3C).

**Figure 3.**
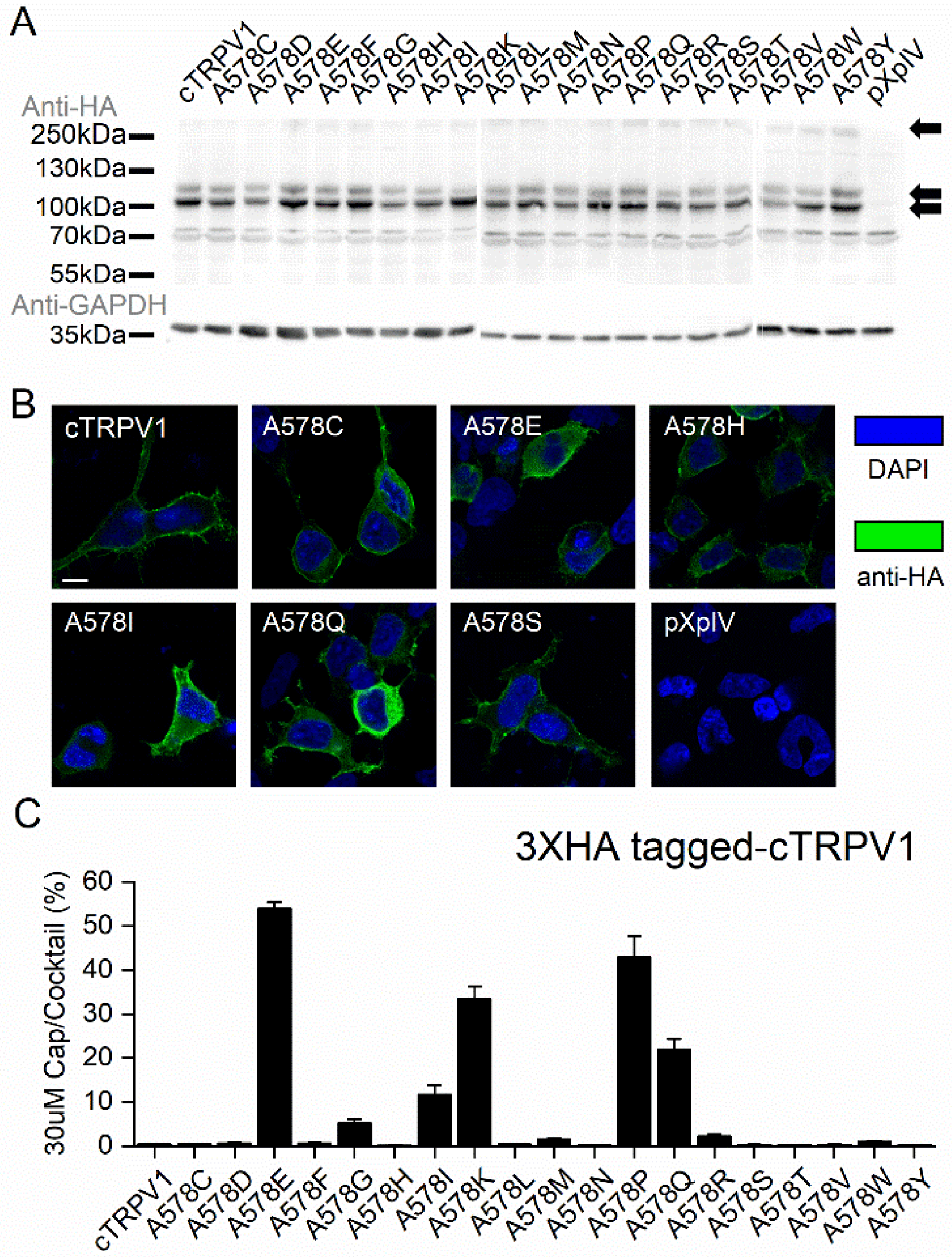
cTRPV1 with A578 residue mutation analysis. (A) Western blotting of cTRPV1 wild type and mutants. (B) Immunostaining of cTRPV1 wild type and mutants expressed on HEK293T (scale bar = 10μm). (C) Capsaicin 30 μM (N=3) induced activation normalized to cocktail (N=2∼3) induced maximum channel opening.

### The hydrophilic derivative of capsaicin activate rTRPV1 and cTRPV1-A578E

A cryo-EM structure with antagonist capsazepine (capZ) bound to rTRPV1 shows the Y511, S512, and E570 sidehains are close enough to interact with its catechol group, which consists of two aromatic hydroxyl groups, compared with a hydroxyl and methoxy group of vanilloids (Gao et al., 2016) (Fig. 4A). To better understand how point mutation on cTRPV1 affect capsaicin interaction with the channel, we synthesized a series of capsaicinoids (Fig. 4B) with altered vanilloid groups. Capsaicin has a polar vanilloid headgroup linked via amide to an aliphatic tail, and functionalization of the vanilloid phenol oxygen has been shown to reduce, but not eliminate, channel activation (Li et al., 2011). Capsaicinoids were prepared by alkylation of the potassium phenolate salts of capsaicin (Materials and Methods) to give hydrophilic *O*-ethylamine (Cap-EA) or *O*-ethylmethylamine (Cap-EMA) derivatives. To determine how the capsaicin interacts with the capsaicin sensitive residues of cTRPV1-578, we tested our capsaicin sensitive mutants with Cap-EA or Cap-EMA (Fig. 4B) to replace the hydroxyl group on vanilloid (Li et al., 2011). To exclude the effect of reduced permeability across plasma membrane, the TRPV1 expressing HEK293T was pre-treated with 10 μM capsaicin derivative dissolved in Ca^2+^-free imaging solution. Upon recording, 10 μM ligand and 2 mM Ca^2+^ in imaging solution was added to elicit extracellular Ca^2+^ entry. For Cap-EA, the pre-incubation time was 5 minutes; and for EMA the pre-incubation time was 1 hour. The induced response was compared with 10 μM capsaicin-induced response that use the same method for pre-treatment and stimulation. Both capsaicin derivatives are sufficient to activate rTRPV1 but not cTRPV1. However, Cap-EA only successfully activates Ca^2+^ current from the cTRPV1-A578E mutant and partially activate the cTRPV1-A578K (Fig. 4C). Cap-EMA slightly activate the cTRVP1-A578K and E (Fig. 4D). cTRPV1-A578P and Q were failed to be activate by the capsaicin derivatives. The replacement of hydroxide group with other hydrophilic side chain drastically reduce the accessible amino acids to activate the channel. Stimulate A578 mutants of cTRPV1 apparently has stringent structural requirements.

**Figure 4.**
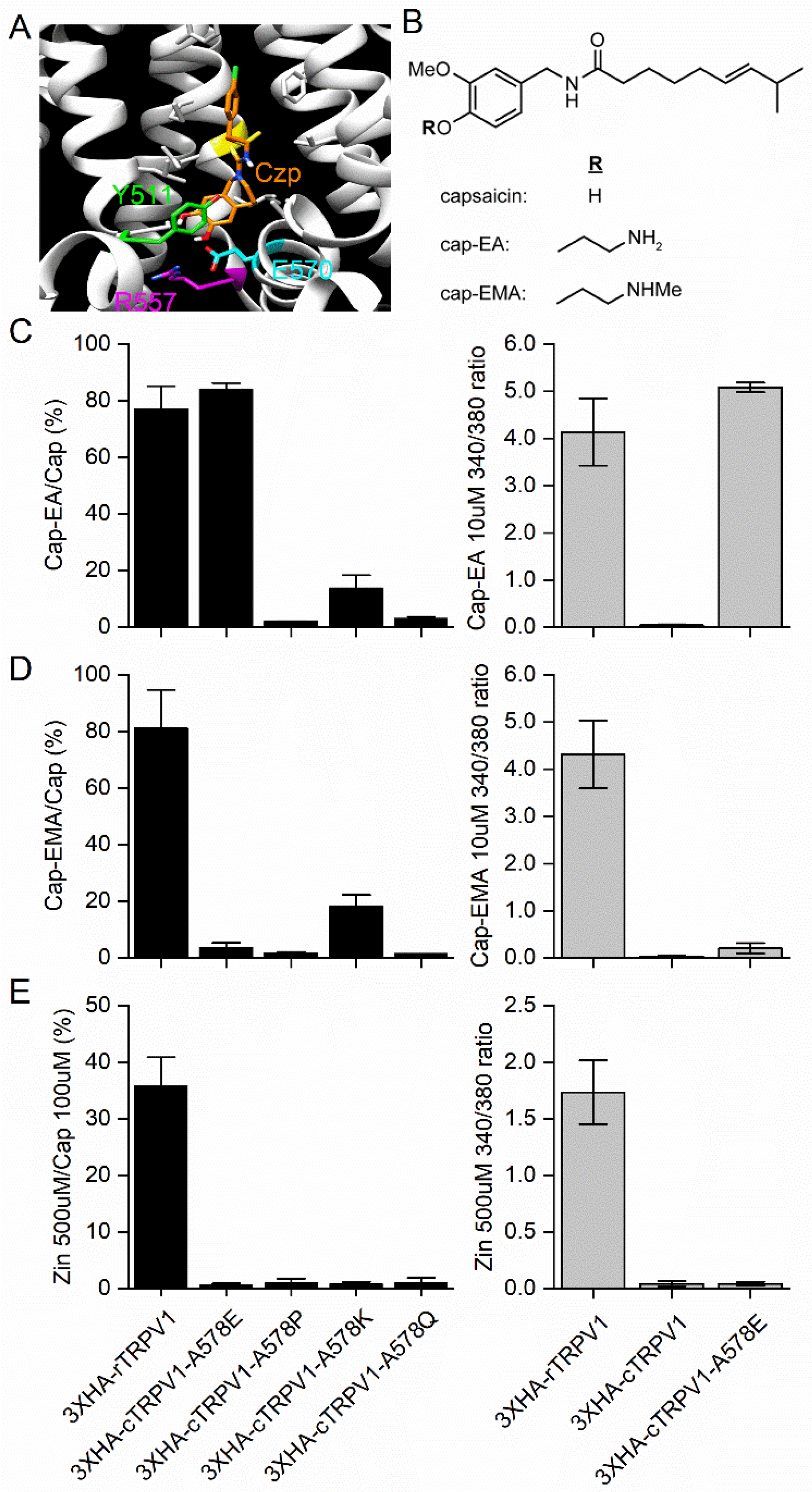
Testing the vanilloids sensitivity of rTRPV1 and cTRPV1-A578 capsaicin active mutants. The right graph shows the 340/380 ratio of rTRPV1, cTRPV1, and cTRPV1-A578E. (A) The Cryo-EM image from PDB 5is0 with some critical residues close to catechol group highlighted with colors. (B) Structures of synthetic capsaicinoids cap-EA and cap-EMA. (C) Cap-EA 10 μM induced activation on rTRPV1 and cTRPV1 capsaicin sensitive mutants normalized to capsaicin 10 μM induced response (N=3). (D) Cap-EMA 10 μM induced activation of rTRPV1 and cTRPV1 capsaicin sensitive mutants (N=3). (E) Zingerone 500 μM induced TRPV1 response was normalized to 100 μM capsaicin response (N=3).

We test the receptors with the natural product zingerone in the ginger to determine if the vanillod group is sufficient to activate the mutants. Zingerone relatively weak vanilloid agonist that need much higher concentration than capsaicin to cause the stimulation (Liu and Simon, 1996). Zingerone lacks the aliphatic tail and acyl-amide moiety presenting on capsaicin. Zingerone 500 μM successfully activates the rTRPV1, but failed to activate cTRPV1-A578 mutants (Fig. 4E). The capsaicin 100 μM positive control activated rTRPV1 and all the capsaicin sensitive mutants, but not wild-type cTRPV1. Our result shows the vanilloid part alone is not strong enough to activate the channel with theoretically functional 578 residues.

### The cTRPV1-A578 and rTRPV1-E570 have similar amino acid preferences for capsaicin

For rat TRPV1, even though all the rTRPV1-E570 mutants protein expression determined by Western blotting (Fig. 5A) and the protein membrane distribution (Fig. 5B) looked similar between mutant and wild type, the cocktail did not successfully activate most of the mutants (data not shown). Therefore, we normalized the capsaicin-induced response to protein level determined by Western blotting (Table S2). Even the protein expression level of mutants showed large variation on Western blotting. rTRPV1 is known to cause obvious cytotoxicity to the transfected cell line, and might cause the protein amount variation by killing the high expression cells. E570K, and Q mutants exhibit responsiveness to 300nM capsaicin as wild type. The E570P decrease the capsaicin sensitivity of rTRPV1 even the protein amount of mutant is similar to wild type (Fig. 5C). We expected the amino acid sharing similar property important for cTRPV1-A578E to sense capsaicin will also respond to capsaicin treatment. Rat and chicken TRPV1 partially share the same preference to specific amino acid to constitute capsaicin sensitivity, but the species specific functional amino acid also exist. The cTRPV1-A578I (Fig. 3C) and rTRPV1-E570R (Fig. 5C) also seem to sense capsaicin for the 340/380 ratio increase after adding the agonist, but the strength is only about 10% compared to the maximum activation of channel or rTRPV1 wildtype. We don’t sure if the small fluorescence ratio value increment has physiological relevance and do not have further discussion.

**Fig 5.**
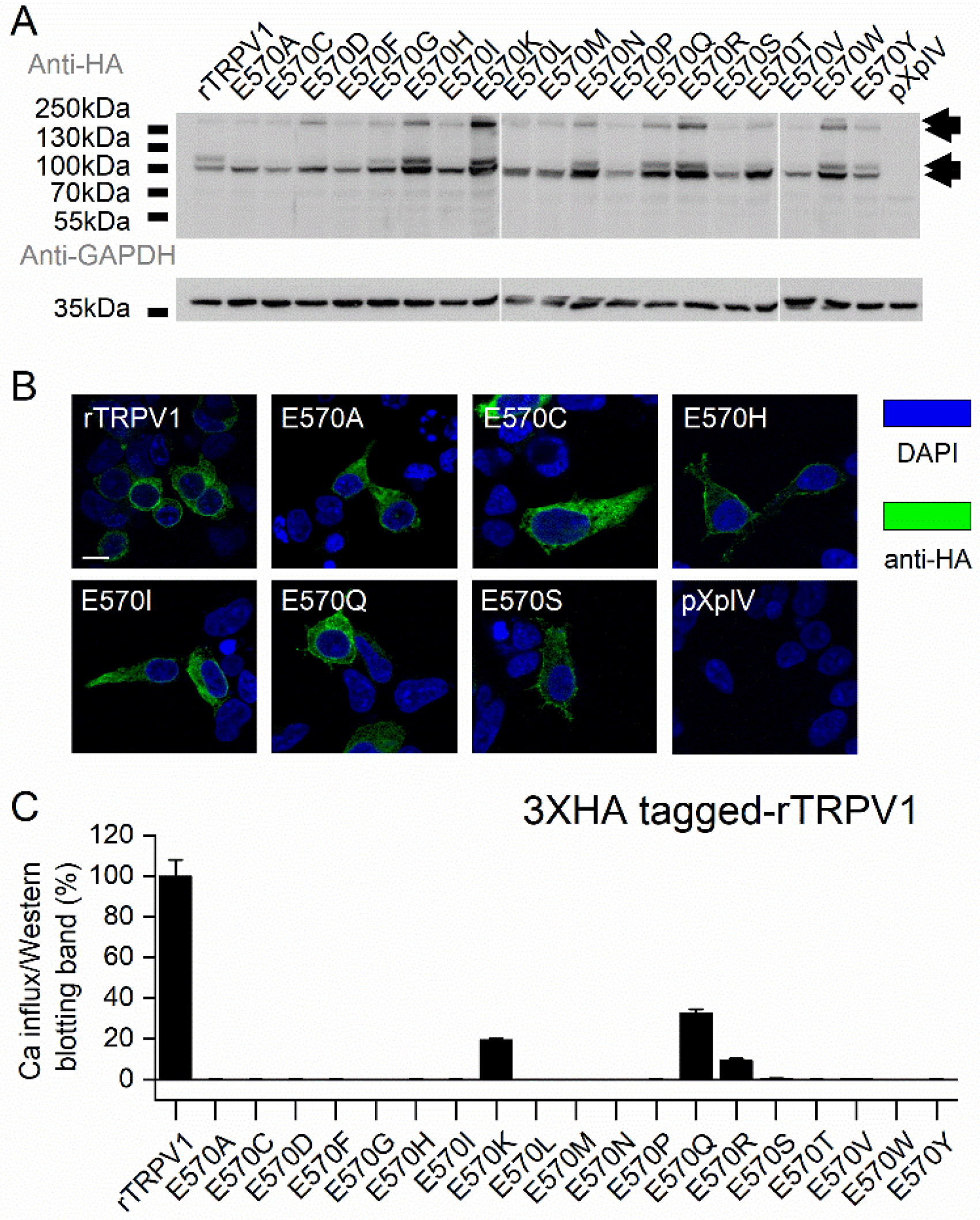
Analysis the corresponding reside mutants on rTRPV1. (A) Western blotting of rTRPV1 wild type and mutants. (B) Immunostaining of rTRPV1 wild type and mutant expressed on HEK293T (scale bar=10μm). (C) Capsaicin 300nM (N=3) induced activation normalized to protein expression level determined by Western blotting.

## Discussion

Animals living in various environments with diverse survival strategy would have to generate specific physiological adaptation. TRPV1 is critical for determining temperature and noxious chemical sensation. Activation of TRPV1 is the principle excitatory mechanism of chemical and thermal nociceptor. TRPV1 is familiar for human to taste spicy food or feel noxious heat. In vampire bat and zebrafish, the TRPV1 orthologues have lower temperature threshold and function as detector to heat source in environments (Gau et al., 2013; Gracheva et al., 2011). Squirrels and camels otherwise have TRPV1 with reduced temperature sensitivity adapting to the hot environment (Laursen et al., 2016). The molecular mechanism of the TRPV1 functional flexibility in different animals had been widely examined to elucidate how to control the channel, which is critical in painful sensation for human.

Chick TRPV1 insensitivity to capsaicin has been attributed to the need for seed dispersal. Capsaicin is reported to selectively discourage packrat and cactus mouse but not thrasher. In contrast to the seed predator rodents, the seeds consumed by thrasher withstand good germination rate (Tewksbury and Nabhan, 2001). It indicates the directed deterrence in chili paper may shape plant-vertebrate interactions. Nonivamide and resinferatoxin in hot peppers are also vanilloids that can activate TRPV1 (Constant et al., 1996; Raisinghani et al., 2005). Avian TRPV1 is relatively insensitive to capsaicin. Thus, chicken can be fed with dried chili pepper and color the egg yolk to orange. Knowing how the TRPV1-expressing nociceptors distinguish different strength and types of stimulation is important for specifically targeting nociceptors working in different analgesic and hyperalgesia issues. It’s theoretically important to understand the vanilloid induced activation in order to generate competitive antagonists, such as capsazapine, 6-iodononivamide, and 6-iodo-RTX.

Chicken TRPV1 is usually classified as capsaicin insensitive, although treatment with high concentrations (100 μM) of capsaicin in this study trigger a small response (Fig. 1D). High sequence similarity between chicken and rat TRPV1 (Jordt and Julius, 2002) suggests that acquiring capsaicin sensitivity depends on minor sequence changes. Testing capsaicin on the newly cloned TRPV1 orthologs is often used to determine the existence of vanilloid sensitivity of these protein. *Xenopus tropicalis*, rabbit, and zebrafish orthologs are classified as weak capsaicin sensors. Corresponding sites of rat S512 and T550 on these species have been shown to play key roles in capsaicin sensitivity by sequence comparison and site-specific mutagenesis (Gau et al., 2013; Gavva et al., 2004; Ohkita et al., 2012).

In this study, we have constructed rat and chicken chimeric TRPV1 channels and point mutants for testing a series of capsaicinoids for their ability to mediate Ca^2+^ influx. The exchanged sites can be classified as four functionally different regions on TRVP1. The TRPV1 transmembrane domain is responsible for ligand sensitivity. The N-, and C-termini play a role in modulating the activation strength and sensitivity (Chuang and Lin, 2009; Lishko et al., 2007). Alternative exon splicing or partially lack of C-terminus amino acid generate TRPV1 channels with lower temperature thresholds (Gracheva et al., 2011). The truncation of large portion in C-terminal generate a non-functional channel (Vlachova et al., 2003). It seems intuitive binding sits for lipophilic ligands are located within the transmembrane domain, and multiple studies have identified critical transmembrane residues as responsible for vanilloid sensitivity in rTRPV1. However, there are only limited reports on cTRPV1, leaving open the possibility that the N and C-termini of rat may mediate species differences. Our result shows the chimeric channel with transmembrane domain have obvious capsaicin sensitivity while rat N or C-terminus do not.

We find that, within the transmembrane region, deletion of G602-N625 impairs both the channel opening and the capsaicin induced activation in our study. The partial deletion of this region was reported to impair the heat sensitivity of mouse TRPV1, and full deletion eliminate channel current (Cui et al., 2012). By A578E mutagenesis on cTRPV1, we proved the single point mutation can make the cTRPV1 become sensitive to capsaicin, and we find the K, Q and P also increase the cTRPV1 capsaicin sensitivity. The conversion, however, is limited by the structure of capsaicin itself, for we test the capsaicin analog with hydrophyilic moiety on the vanilloid head generate restriction for the amino acid types and decrease the potency of drug on rTRPV1. Only the cTRPV1-A578E is strongly activated by Cap-EA (ethylamine) but not Cap-EMA (methylethylamine). cTRPV1-A578K is slightly activated by both analogs. We further confirm the conversion is reversible on rat, for the E570A mutation decrease the capsaicin induced response on rat TRPV1. Our finding is in agreement to the importance of E570 in rat TRPV1 for sensing capsaicin. Yang et. al. (2015) shows that the E570 (E571 on the paper) is responsible to form hydrogen bond with the vanillyl head of capsaicin to pull the S4-S5 linker and make the channel pore open. The mutation not only decrease the capsaicin sensitivity of murine channel but also make the channel have smaller open probability for capsaicin. It shows the residue on the rat TRPV1 is critical for capsaicin binding and channel gating. The systematic mutagenesis on rTRPV1 shows the E570Q and K also possess capsaicin sensitivity. We tested the mutant sensitivity with zingerone, which has a short aliphatic tail, and find that without van der Waals interactions between aliphatic tail and binding pocket, a high concentration is required to activate even the rTRPV1. All the capsaicin-sensitive cTRPV1 mutants do not respond to 500 μM zingerone, and suggest the aliphatic tail of vanilloid is also an important feature to activate the cTRPV1. Both results suggesting the residue on rat and chicken TRPV1 share similar mechanisms for capsaicin-mediated channel opening. The structural requirements of the cTRPV-A578 mutant to sense capsaicinoids implies that the mutation participate in the capsaicin binding. Although we take the advantage of using cocktail containing multiple strong TRPV1 agonist to prove the mutated channel is functionally active or not, we unexpectedly face the problem of activating the rTRPV1 with mutation on E570. The residue on S4-S5 is responsible to transfer the capsaicin binding induced conformational change to S5-S6 pore domain. We would expect the the PAO potentiated rTRPV-E570 mutants still possess some vanilloid sensitivity and can be activated. The severe impair of channel to open in response to cocktail shows the residue can also cause some other unknown limitation to the channel functionality.

The approach we report here to expressed mutated cTRPV1 on the A578 residue critical for capsaicin-induced activation allow us to conclude the vanilloid sensitivity difference between avian and mammal owe to one amino acid difference. The alanine at position 578 residue is conserved between some avian orthologues TRPV1 sequences (e.g., duck, XM_021274874; ostrich, XM_009672965; hummingbird, XM_008495099). The residue is glutamic acid on *Xenopus tropicalis* and snake, and glutamine on zebrafish. It shows this mechanism of loss of capsaicin sensitivity is emerge in the avian. Although partially different amino acids preference reduces the analogy of vanilloid binding process between rat and chicken TRPV1, it proves more flexibility to explain the mechanism for capsaicin-induced activation. In any event, the result shows the possibility to constitute capsaicin sensitivity by adding one mutation.

## List of Symbols and Abbreviations

TRPV1: transient receptor potential cation channel subfamily V member 1
Cap: capsaicin
RTX: resiniferatoxin
RR: ruthenium red
PAO: phenylarsine oxide
Cap-EA: *O*-aminoethyl capsaicin
Cap-EMA: *O*-(*N*-methylaminoethyl) capsaicin
Zin: zingerone

---

**Table S1.** Cocktail and capsaicin induced cTRPV1 wild type and mutants activation, present as 340/380 ratio.

|  |  |  |
| --- | --- | --- |
| Gene | Maximum 340/380 ratio-BG<br>(N=1) |  |
| | Cocktail | Cap 30 μM |
| 3XHA-cTRPV1 | 12.03 | 0.08 |
| 3XHA-cTRPV1-A578C | 11.72 | 0.02 |
| 3XHA-cTRPV1-A578D | 3.44 | 0.03 |
| 3XHA-cTRPV1-A578E | 11.29 | 5.77 |
| 3XHA-cTRPV1-A578F | 2.41 | 0.02 |
| 3XHA-cTRPV1-A578G | 2.68 | 0.17 |
| 3XHA-cTRPV1-A578H | 14.14 | 0.02 |
| 3XHA-cTRPV1-A578I | 11.46 | 0.76 |
| 3XHA-cTRPV1-A578K | 12.60 | 4.36 |
| 3XHA-cTRPV1-A578L | 8.23 | 0.03 |
| 3XHA-cTRPV1-A578M | 11.34 | 0.15 |
| 3XHA-cTRPV1-A578N | 7.42 | 0.02 |
| 3XHA-cTRPV1-A578P | 6.51 | 3.84 |
| 3XHA-cTRPV1-A578Q | 13.05 | 2.93 |
| 3XHA-cTRPV1-A578R | 13.34 | 0.17 |
| 3XHA-cTRPV1-A578S | 13.98 | 0.06 |
| 3XHA-cTRPV1-A578T | 14.62 | 0.02 |
| 3XHA-cTRPV1-A578V | 10.81 | 0.04 |
| 3XHA-cTRPV1-A578W | 2.21 | 0.02 |
| 3XHA-cTRPV1-A578Y | 9.17 | 0.02 |

**Table S2.** Quantitative result of rTRPV1 wild type and mutant Western blotting band intensity.

|  |  |  |  |  |  |
| --- | --- | --- | --- | --- | --- |
|  | Western 1 | Western 2 | Western 3 | Average | s.e.m |
| WT | 1 | 1 | 1 | 1 | 0 |
| A | 0.52 | 1.01 | 0.87 | 0.80 | 0.15 |
| C | 1.04 | 0.61 | 0.74 | 0.80 | 0.13 |
| D | 1.01 | 1.86 | 1.36 | 1.41 | 0.25 |
| F | 1.10 | 1.09 | 0.58 | 0.92 | 0.17 |
| G | 1.30 | 2.46 | 0.98 | 1.58 | 0.45 |
| H | 1.16 | 6.19 | 1.54 | 2.96 | 1.62 |
| I | 0.21 | 2.15 | 0.50 | 0.95 | 0.61 |
| K | 0.99 | 7.59 | 1.77 | 3.45 | 2.08 |
| L | 2.02 | 1.09 | 0.77 | 1.29 | 0.37 |
| M | 1.86 | 1.17 | 0.68 | 1.24 | 0.34 |
| N | 4.39 | 2.78 | 1.13 | 2.77 | 0.94 |
| P | 1.23 | 0.72 | 0.41 | 0.78 | 0.24 |
| Q | 6.38 | 2.61 | 0.95 | 3.31 | 1.61 |
| R | 5.07 | 3.65 | 1.51 | 3.41 | 1.03 |
| S | 0.95 | 0.92 | 0.77 | 0.88 | 0.06 |
| T | 1.61 | 2.27 | 0.83 | 1.57 | 0.42 |
| V | 1.59 | 0.68 | 0.53 | 0.93 | 0.33 |
| W | 2.61 | 2.80 | 1.53 | 2.31 | 0.39 |
| Y | 3.43 | 1.23 | 0.60 | 1.75 | 0.86 |

\*Each band intensity was divided to the rTRPV1 band on the same PVDF membrane to reduce the variation caused by photography error.

## Acknowledgements

This project was supported by a MOST grant and a Career Development Award to HC by Academia Sinica. The authors thank colleagues for multiple helpful suggestions in organization till the finished form.

